# Continental-scale microbiome study reveals different environmental characteristics determining microbial richness and composition/quantity in hotel rooms

**DOI:** 10.1101/849430

**Authors:** Xi Fu, Yanling Li, Qianqian Yuan, Gui-hong Cai, Yiqun Deng, Xin Zhang, Dan Norbäck, Yu Sun

**Affiliations:** Guangdong Provincial Key Laboratory of Protein Function and Regulation in Agricultural Organisms, College of Life Sciences, South China Agricultural University, Guangzhou, Guangdong, 510642, PR China; Key Laboratory of Zoonosis of Ministry of Agriculture and Rural Affairs, South China Agricultural University, Guangzhou, Guangdong, 510642, PR China; School of Public Health, Sun Yat-sen University, Guangzhou, PR China; Occupational and Environmental Medicine, Dept. of Medical Science, University Hospital, Uppsala University, 75237 Uppsala, Sweden; Institute of Environmental Science, Shanxi University, Taiyuan, PR China

## Abstract

Culture-independent microbiome surveys have been conducted in homes, hospitals, schools, kindergartens and vehicles for public transport, revealing diverse microbial distributions in built environments. However, microbiome surveys have not been conducted in hotel environments; thus, the composition and associated environmental factors are not clear. We presented the first continental-scale microbiome study of hotel rooms (n = 68) spanning large geographic regions. Bacterial and fungal communities were described by amplicon sequencing of the 16S rRNA gene and internal transcribed spacer (ITS) region and quantitative PCR. Similar numbers of bacterial (4,344) and fungal (4,555) operational taxonomic units were identified, but fungal taxa showed a local distribution compared with bacterial taxa. Aerobic, ubiquitous bacteria dominated the hotel microbiome with high compositional similarity to previous samples from building and human nasopharynx environments. The abundance of *Aspergillus* was negatively correlated with latitude and accounted for ∼80% of the total fungal load in seven low-latitude hotels. We calculated the association between hotel microbial dynamics and 16 indoor and outdoor environmental characteristics. Fungal β-diversity and quantity showed concordant variation and were associated with the same environmental characteristics, including latitude, quality of the interior, proximity to the sea and visible mold, while α-diversity decreased with heavy traffic (95% CI: −127.05 to −0.25) and wall-to-wall carpet (95% CI: −47.60 to −3.82). Bacterial β-diversity was associated with latitude, quality of the interior and floor type, while α-diversity decreased with recent decoration (95% CI −179.00 to −44.55) and mechanical ventilation (95% CI: −136.71 to −5.12).

**Importance:** This is the first microbiome study to characterize microbial composition and associated environmental characteristics. In this study, we found concordant variation between microbial β-diversity and absolute quantity and discordant variation between β-diversity/quantity and α-diversity. Our study can be used to promote hotel hygiene standards and provide resource information for future microbiome and exposure studies associated with health effects in hotel rooms.

## Introduction

Recent advances in culture-free high-throughput sequencing techniques and bioinformatics analyses have greatly facilitated microbiome research in many fields, including human gut, skin and respiratory tract microbiomes and disease, environmental microbiomes, and large-scale communal projects, such as the earth’s microbiome project [1-6]. Although the total number of microbiome studies has increases dramatically in the past few years, they are unevenly distributed among areas. For example, more than 60% of the microbiome studies are restricted to the human gut and skin and laboratory-based model organisms, and only ∼2% of the studies were performed in the built environment [7], which is one of the most important environments in the lives of modern humans. The National Human Activity Pattern Survey from the United States reported that people spend an average 87% of their time in buildings and another 6% of their time using transportation [8]. Furthermore, indoor microbial exposures have been reported to relate to occupant health [9]. Several risk and protective species have been identified to be associated with human noninfectious diseases such as asthma, allergy and respiratory symptoms [10-12], and exposure to high bacterial and fungal diversity has been reported to have protective effects on childhood asthma [13, 14]. A long term goal of indoor microbiome research is to identify a “healthy building microbiome” and promote human well-being, and the necessary first step is to characterize the microbiome composition of different indoor environments.

Indoor microbiome research is a complex multidisciplinary field that requires knowledge of microbiology, ecology, environmental science, building science, and epidemiology while integrating new techniques, such as next-generation sequencing (NGS) [15]. To date, indoor microbiome studies have mainly focused on home environments [16-20], and studies from other environments have also been reported, such as hospitals [21, 22], schools [23], university dormitories [24], kindergarten [23] and vehicles for public transport [25, 26]. In principle, the framework of indoor microbiome, especially in the home environment, has been generally established. Indoor microbes originate from multiple sources, including outdoor air; soil; plants; the human skin, gut and mouth; pets; plumbing systems [27]. It is generally accepted that the outdoor environment and indoor occupants and animals are the two major sources of indoor bacteria [17, 28], but the relative contributions of the two sources vary with building type and location. For example, the percentage of indoor bacteria from human sources varied from 4% in a conference room [29] to >30% in a university housing complex [30]. In a noncontaminated or moldy environment, indoor fungi are mainly sourced from outdoor air and thus structured by climate and geographical patterns [17, 20]. Other indoor factors, such as mechanical ventilation, type of carpet, and cleaning procedure and frequency, also shape indoor fungal communities [10]. Identifying the environmental factors associated with microbial composition promotes the understanding of microbiome dynamics in indoor ecological niches.

Hotels are a common public environments for guests and hotel staff. Each year, millions of guests and travelers stay in hotels, and there is a public health concern regarding hotels’ hygiene standards and practice. Unlike household residences, each hotel room is shared by many guests, but many influencing factors are also controlled. For example, most hotels use standard cleaning procedures and ventilation systems for air exchange, and no pets are allowed in hotel rooms so a major source of indoor microbiome is also controlled. Therefore, hotel rooms are an appropriate place to conduct a global indoor microbiome comparison. As no microbiome survey has been conducted in the hotel environment, it is unclear if the composition and environmental drivers of hotel microbiome are similar to the home environment or not. A few studies have quantified hotel bacterial and fungal taxa by counting colony-forming microbes on medium [31, 32], but due to technique limitations, this approach can identify only less than 1% of total microbial species [33]. Our previous study used qPCR to monitor indoor fungal quantity in European and Asian hotels and identified several associated indoor or outdoor characteristics [34]. A few hotel epidemiological studies have focused on single infectious microbial exposure or outbreaks in hotel rooms, such as *Legionella* and Norovirus infection [35, 36]. Thus, the overall assemblage and diversity of hotel microbes are still largely unknown.

Microbiome study must quantify and disentangle thousands of phylogenetically related or distinct species. Amplicon and shotgun metagenomic sequencing provides relative abundance information for all the microbes in a sample, and α- and β-diversity are statistics commonly used to describe microbial composition and distribution. α-diversity, including the number of observed species, Chao1 index and Shannon index, quantifies the community richness or evenness within an individual sample [37-39]. β-diversity, including quantitative metrics, such as the Bray-Curtis or UniFrac distance, evaluates community differences between samples [40, 41]. Together with quantitative approaches, such as qPCR, the absolute number of microbial cells per taxon can be identified [42], which can be further used to identify the key microbes associated with health outcomes and environmental characteristics. This approach has been applied in several human intestine and indoor microbiome studies [14, 18, 43], and it has been shown to be a solid and powerful approach for characterizing microbiome dynamics and associated factors.

In this study, we sequenced microbial amplicon regions, including the bacterial 16S ribosomal RNA gene and fungal internal transcribed spacer (ITS) region, to characterize the microbial composition of hotel dust in a large geographic area covering 19 European and Asian countries. In total, 16 environmental factors were analyzed together with microbiome data in multiple linear regression and permutation models to identify factors associated with microbial α- and β-diversity and fungal quantity. We further discussed the results and implications from the microbial, ecological and indoor health perspectives.

## Results

### Microbial diversity and composition in hotel dust

In this study, dust swab samples were collected by a professional hygienist from 68 hotels in 19 European and Asian countries. The sampling location was rarely cleaned by hotel staff and thus reflected the airborne microbiome composition accumulated over at least several months. In each hotel, two dust samples were collected in one room from the upper half of the door frame with a swabbing area of 1 x 60 cm. One sample was used for quantitative analysis of fungi DNA, the results of which were published in a previous paper [34], and the other sample was used for 16S rRNA gene and ITS region sequencing in this study. More than 30 environmental characteristics, including latitude, surrounding traffic, quality of the interior (textiles, walls and furniture), and visible mold, were assessed at each sampling site. Pearson’s correlation was conducted to reduce collinearity (ρ > 0.7), and 16 environmental characteristics were kept for further analysis (Table 1).

**Table 1.**
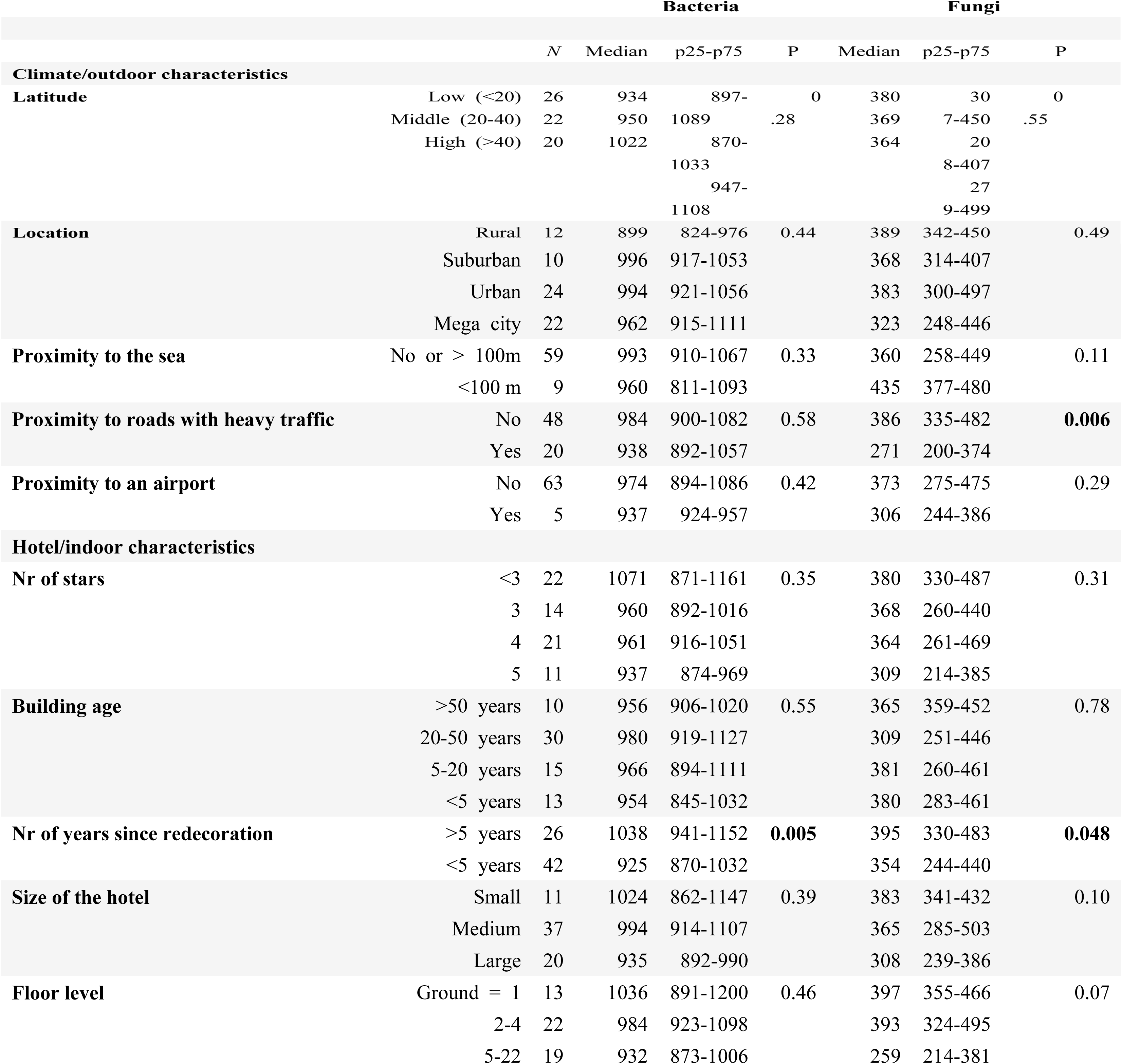
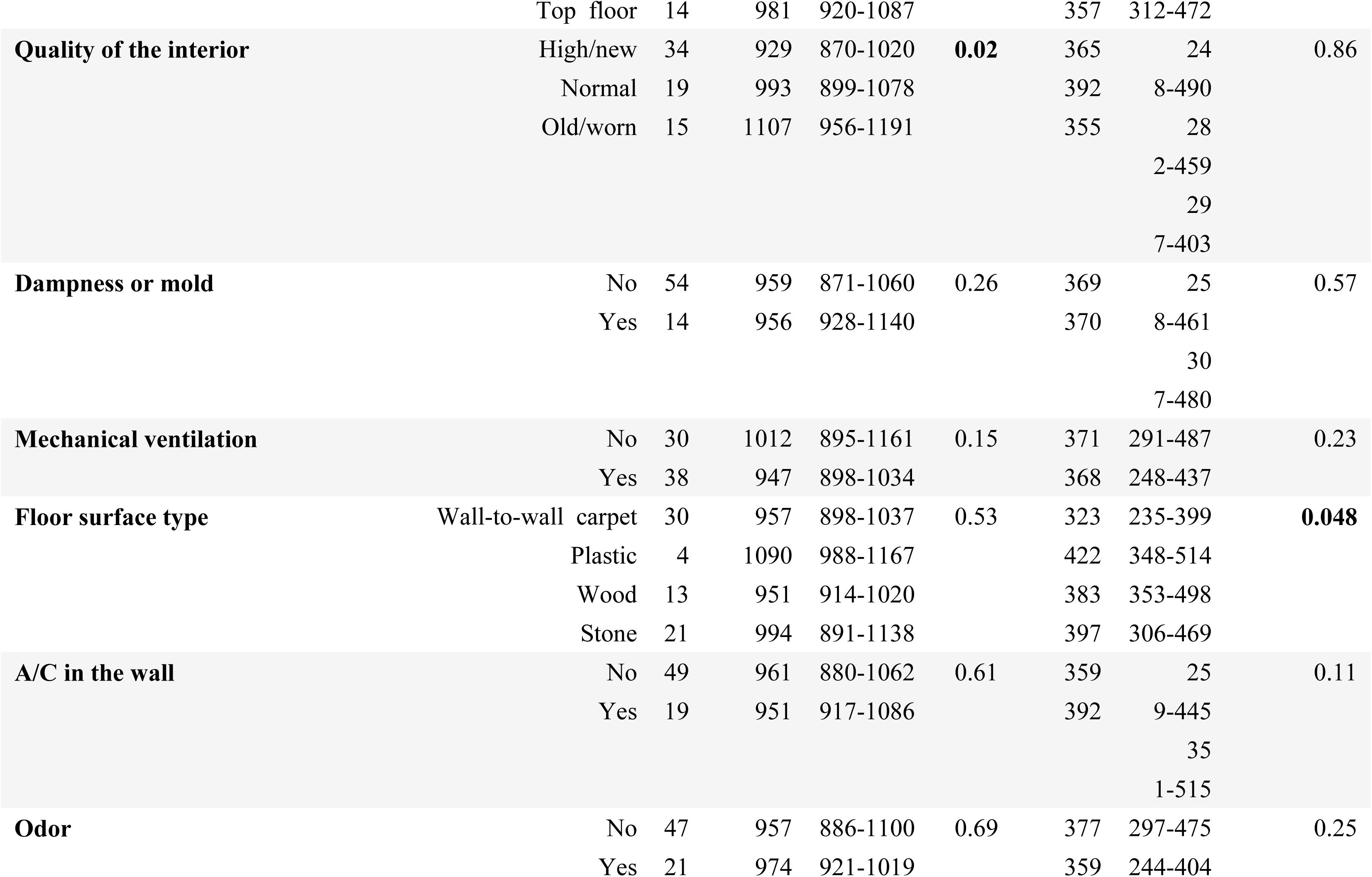
Association between outdoor/indoor characteristics and observed microbial OTUs. P values were calculated by a Kruskal-Wallis test, and p values less than 0.05 are formatted with bold font.

Based on gel electrophoresis of the negative control (Figure S1 and S2), no clear DNA contamination in the reagents or amplification processes was observed for 16S and ITS sequencing. For the sequence data, several quality-control and filtration steps were conducted (see Methods), and rarefaction analysis indicated that the sequencing depth was sufficient to cover the majority of the microbial diversity (Figure S3 and S4). A total of 4,344 bacterial and 4,555 fungal OTUs were obtained, and each dust sample harbored 988 bacterial and 370 fungal OTUs on average. Bacterial and fungal samples had distinct OTU distribution patterns (Figure 1A, 1B). The fungal OTUs were locally distributed; approximately half of the OTUs were found in 1 or 2 samples, and only 1 OTU (*Candida albicans*) was found in all samples. The bacterial OTUs were more widespread on a regional or continental scale; more than half of the OTUs were found in 10 or more samples and 43 OTUs were found in all samples.

**Figure 1.**
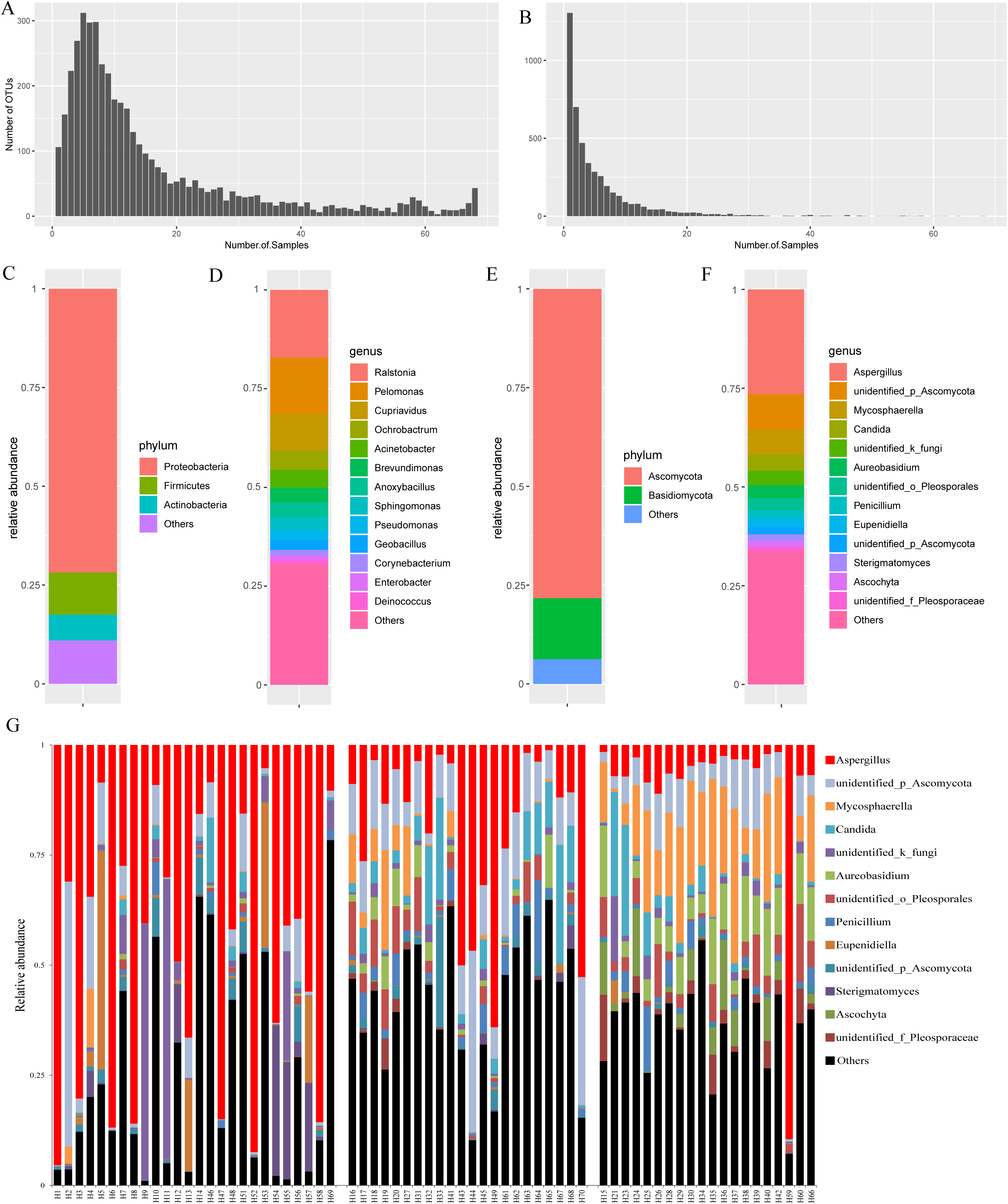
Distribution and relative abundance of bacterial and fungal taxa in hotels. Frequency spectrum for (A) bacterial and (B) fungal OTUs in all samples. The average relative abundance of bacteria at the phylum (C) and genus levels (D). The average relative abundance of fungi at the phylum (E) and genus levels (F). Relative abundance of fungi at the genus level for all samples (G).

Taxonomic information is presented at the phylum and genus levels based on search results from Silva database (Figure 1 C-F). For bacteria, the dominant phylum was Proteobacteria (mean ± standard deviation, 71.8% ± 11.2%), followed by Firmicutes (10.7% ± 4.2%) and Actinobacteria (6.5% ± 6.4%) (Figure 1C, S3 and Table S1). The top genera were the environmental bacteria *Ralstonia* and *Pelomonas*, with an average abundance higher than 10%, followed by *Cupriavidus, Ochrobactrum, Acinetobacter, Brevundimonas, Anoxybacillus, Sphingomonas, Pseudomonas, Geobacillus* and *Corynebacterium* (Figure 1D, S4 and Table S2). We conducted bacterial microbiome novelty score analyses on the Microbiome Search Engine by calculating the compositional similarity between the hotel samples and over 170,000 curated microbiome samples [44]. The microbiome novelty scores were higher than 0.12 for 70% of the hotel samples, indicating high novelty and low compositional similarity to previously published microbial samples (Table S3). The most similar samples were mainly from building and human nasopharynx and skin environments.

The dominant fungal phylum was Ascomycota (78.3% ± 13.8%), followed by Basidiomycota (15.4% ± 11.1%; Figure 1E, S5 and Table S4). The most abundant genus was *Aspergillus*, with an average abundance higher than 25%, followed by *Mycosphaerella, Candida, Aureobasidium* and several unidentified Ascomycota (Figure 1F, S6 and Table S5). The top fungal genus *Aspergillus* had an uneven distribution among samples, with an increasing trend from high to low latitude (Figure 1G). The relative abundance of *Aspergillus* was over 80% in eight hotels, seven of which were from low latitudes, including four in Malaysia, two in Thailand, and one in Vietnam. For the high-latitude hotels, the abundance of *Aspergillus* was lower than 15%, except in Venice (Figure 1G).

The variation in microbial composition was visualized by principal coordinates analysis (PCoA) based on a weighted UniFrac distance matrix for bacteria and a Bray-Curtis dissimilarity matrix for fungi (Figure 2). The high-, middle- and low-latitude samples were generally separated along axes 1 and 2, suggesting that latitude is an important factor structuring the microbiome composition variation in hotel dust. The arch distortion [45] for fungi suggests that axis 1 is a single dominant factor, whereas axis 2 plays a rather minor role in structuring community variation.

**Figure 2.**
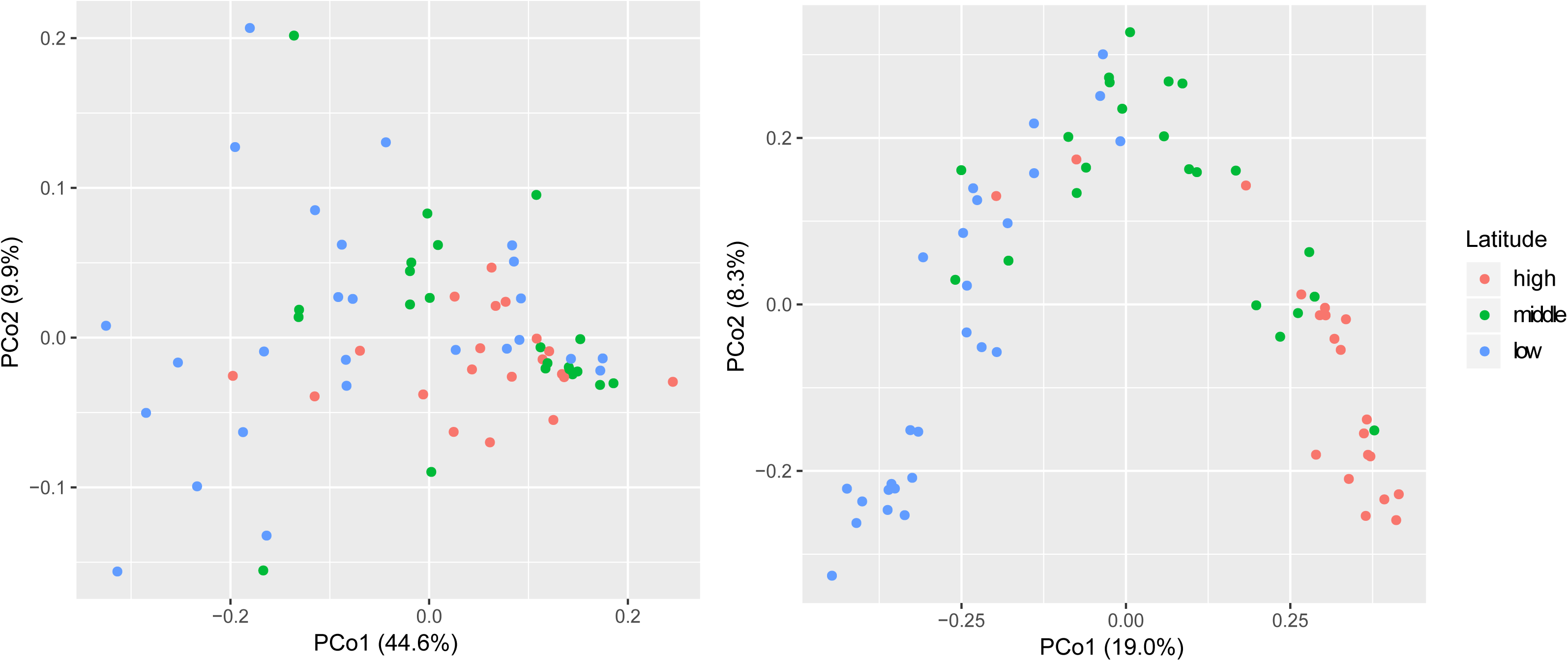
Principle coordinate analyses (PCoA) of (A) bacterial and (B) fungal composition at different latitudes. The PCoAs were based on a weighted UniFrac distance matrix for bacteria and a Bray-Curtis dissimilarity matrix for fungi

### Bivariate analyses between environmental and microbiome characteristics

As a next step, associations were analyzed between environmental factors and microbial α- and β-diversity and fungal DNA quantity (Table 1). The number of observed OTUs was calculated from 27,000 rarefied reads for all samples. For bacteria, recent redecoration and quality of the interior were associated with a decreased number of observed OTUs in dust samples (p=0.005 and 0.02, respectively; Kruskal-Wallis test). For fungi, proximity to roads with heavy traffic, recent redecoration, and the floor surface type of wall-to-wall carpet were associated with a decrease in species richness (p=0.006, 0.048 and 0.048, respectively).

Latitude is an important factor shaping bacterial and fungal community variation (p < 0.001, R^2^ = 0.10 and 0.19, respectively; Adonis analysis; Table S6), consistent with the PCoA results. Other factors, such as floor surface type, seaside location, floor level, quality of the interior, mechanical ventilation and air conditioner in the wall, were also associated with bacterial or fungal β-diversity.

Our previous study used two sets of primers to quantify absolute fungal DNA in these hotel rooms [34]. One primer targeted the fungal ITS1 region and captured a wide range of indoor fungi (>530 species), and the other primer targeted fungal 28S rRNA and captured mainly *Aspergillus* and *Penicillium* taxa (>140 species) [46]. We named the two fungal quantifications fungal DNA 1 and fungal DNA 2 (Table S7 and Table 2), respectively, and analyzed the association with environmental characteristics by a Kruskal -Wallis test. Latitude, proximity to the sea, quality of the interior, dampness or mold, and mechanical ventilation were associated with fungal DNA variation for both datasets (p < 0.05, Table S8).

**Table 2.**
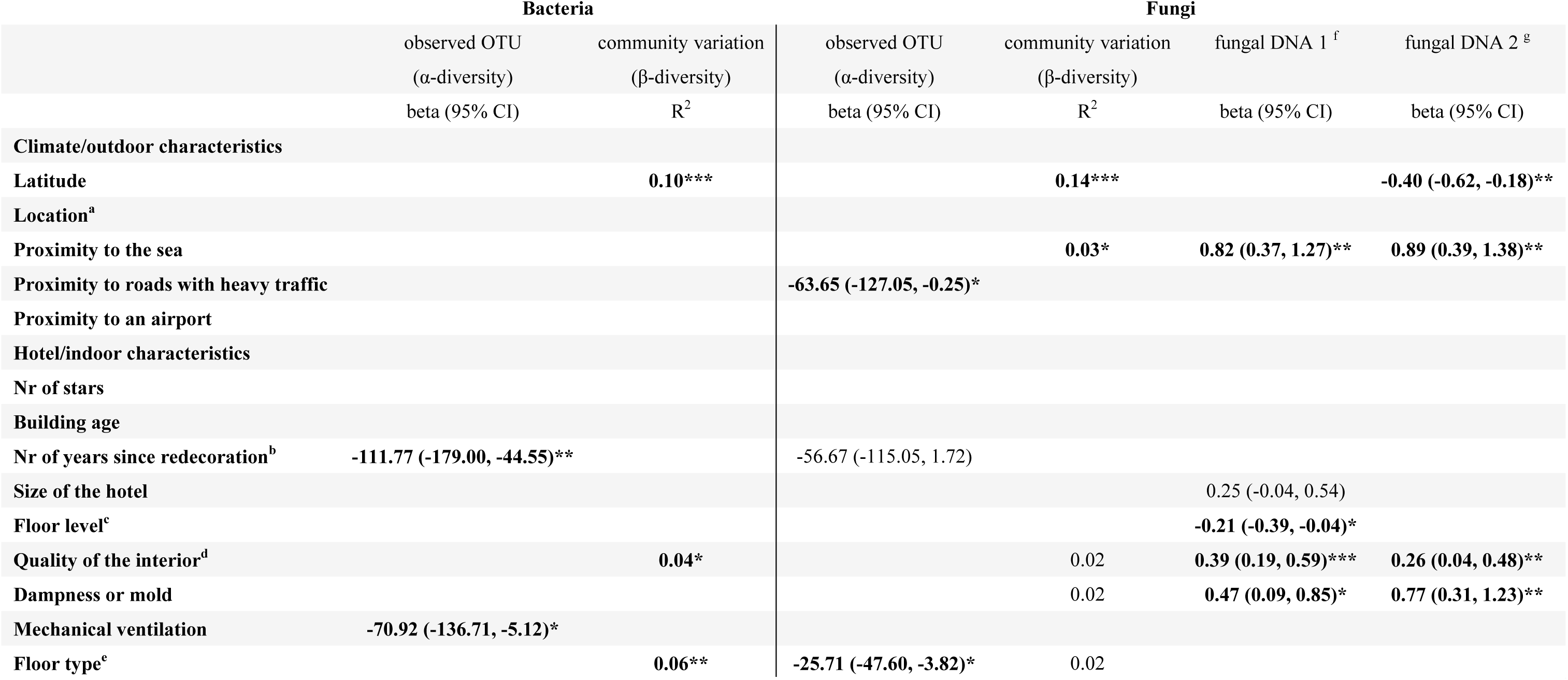
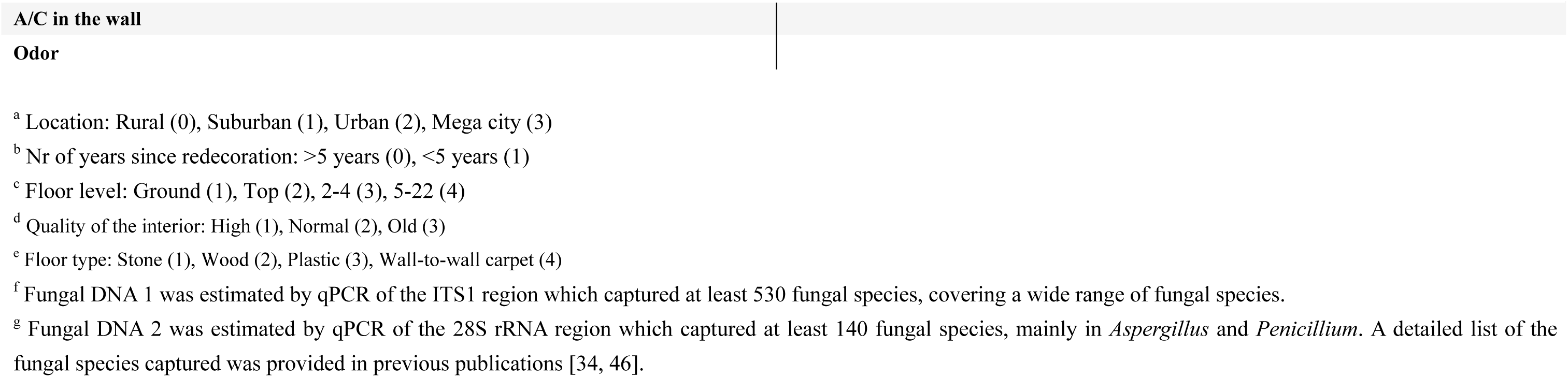
Multivariate analysis between outdoor/indoor characteristics and microbial α- and β-diversity and fungal quantity. The α-diversity and fungal quantity were calculated by forward stepwise linear multiple regression, and β-diversity was calculated by forward stepwise Adonis multivariate analysis with 10,000 permutations. Associations with p values less than 0.05 are formatted with bold font and stars (*** p < 0.001, ** p < 0.01, * p < 0.05), and associations with 0.05 < p < 0.1 are presented without bold font.

### Multivariate analyses between environmental and microbial characteristics

We conducted forward stepwise linear multivariate regression and Adonis analysis between environmental and microbial characteristics (Table 3). The environmental characteristics with a p value < 0.2 in the bivariate analysis were included in the multivariate analyses. Latitude was the strongest factor associated with bacterial community variation (p < 0.001, R^2^ = 0.10), followed by floor type and quality of the interior (p < 0.05, R^2^ = 0.06 and 0.04, respectively). Recent redecoration and the presence of mechanical ventilation decreased bacterial richness in the hotel rooms (beta=-111.77, 95% CI: −170.00 to −44.55; beta=-70.92, 95% CI: −136.71 to −5.12). Thus, bacterial α- and β-diversity were associated with different environmental factors.

Latitude was also the strongest predictor of fungal community variation (p < 0.001, R^2^ = 0.14, Adonis analysis), and was significantly associated with fungal quantity (regression coefficient beta = −0.40, 95% confident interval −0.62 −0.18). Similarly, proximity to the sea, low quality of the interior and visible mold were significantly or marginally significantly associated with fungal community variation (p = 0.03, 0.07 and 0.07 and R^2^=0.03, 0.02 and 0.02, respectively), and all these factors significantly increased fungal DNA quantity (beta=0.82, 95% CI: 0.37 to 1.27; beta=0.39, 95% CI: 0.19 to 0.59; and beta=0.47, 95% CI: 0.09, 0.85, respectively) but did not affect fungal α-diversity. Two factors were significantly associated with fungal α-diversity. The floor type of wall-to-wall carpet decreased fungal richness (beta=-25.71, 95% CI: −47.60 to −3.82). Proximity to roads with heavy traffic decreased fungal α-diversity (beta=-63.65, 95% CI: −127.05 to −0.25). Thus, the factors associated with fungal β-diversity were also associated with fungal absolute quantity, while microbial α-diversity was affected by other environmental factors.

### Environmental factors and associated microbes

We further characterized and visualized bacterial and fungal genera associated with different environmental factors, including latitude, quality of the interior and visible mold. We first plotted taxa with more than 1% variation in abundance between latitudes (Figure 3). For bacteria, *Ralstonia* and *Pelomonas* were significantly more abundant at high latitudes (Kruskal-Wallis test, p < 0.05), while *Cupriavidus* and *Saccharopolyspora* were more abundant at low latitudes (p < 0.05). For fungi, *Aspergillus, Eupenidiella, Sterigmatomycetes, Devriesia, Schizophyllum* and *Exobasidium* were more abundant at low latitudes (p < 0.05), while *Mycosphaerella, Aureobasidium, Penicillium, Malassezia, Cryptococcus, Simplicillium, Botrytis, Stemphylium, Ascochyta* and two unidentified genera were more abundant at middle or high latitudes (p < 0.05).

**Figure 3.**
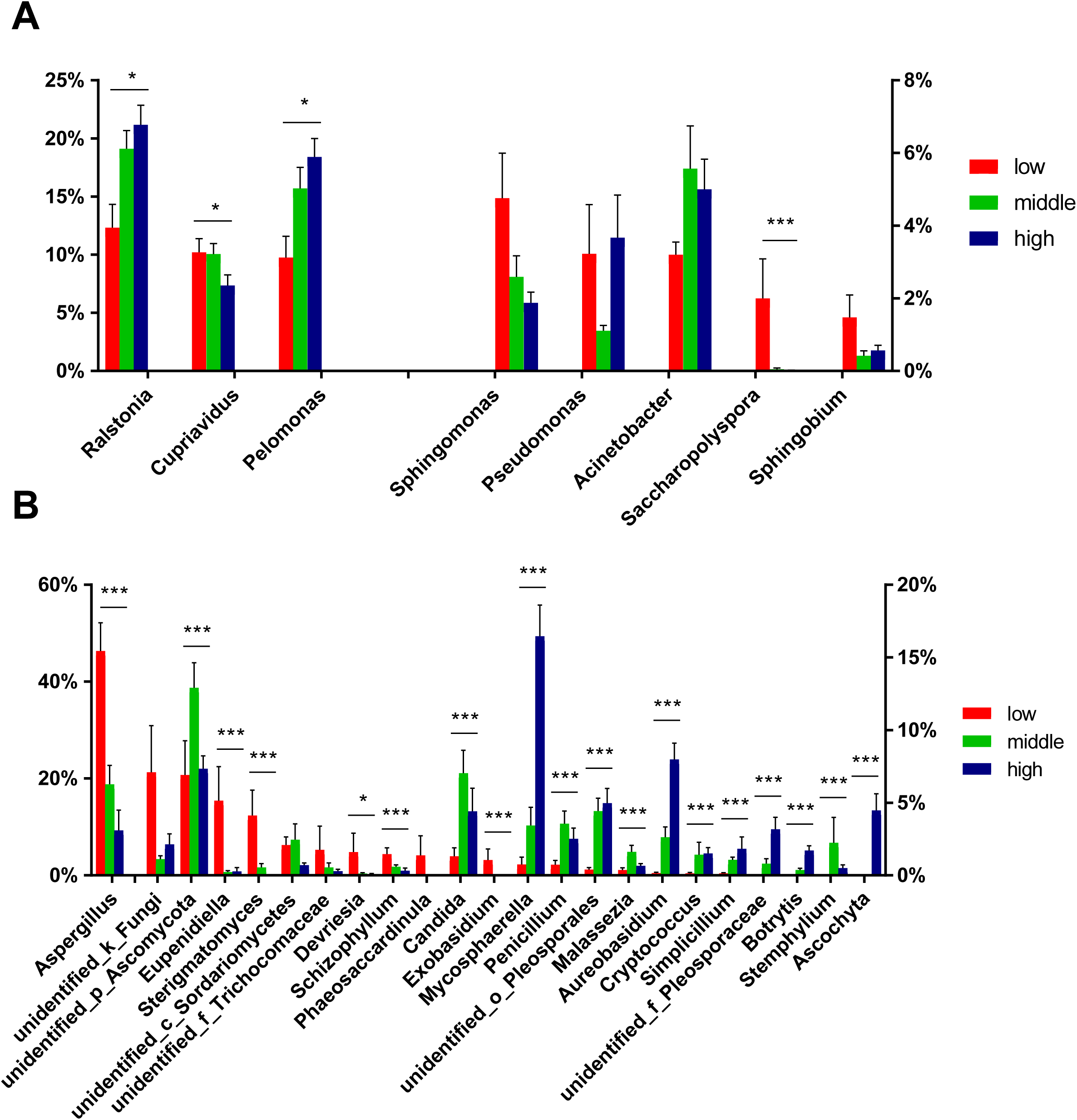
Relative abundance of (A) bacterial and (B) fungal genera at different latitudes. Only genera with relative abundance differences > 1% among latitudes are plotted. Error bars represent the standard error, and a Kruskal Wallis test was conducted (*** p < 0.001, ** p < 0.01, * p < 0.05).

*Saccharopolyspora* was the only bacterial genus associated with the quality of the room interior (textiles, walls and furniture), and the genus was more abundant in worn and old rooms (Figure S8). The relative abundance of *Aspergillus* was 54.9% in rooms with visible mold or dampness but only 19.2% in rooms without mold (p < 0.001; Figure S9), which is consistent with the fact that *Aspergillus* is a common mold in the indoor environment. *Eupenidiella* and *Exobasidium* were also more abundant in moldy hotel rooms (p < 0.05; Figure S9).

## Discussion

In this study, we conducted amplicon sequencing to identify the microbiome community in hotels in 19 countries and the associated environmental characteristics. We found that latitude, seaside location, quality of the interior, and visible mold were associated with both fungal β-diversity and absolute quantity, whereas fungal richness was associated with other environmental characteristics. Similarly, bacterial β-diversity and α-diversity were also affected by different sets of environmental characteristics.

Several studies have characterized microbial α- and β-diversity and quantity in built environments. A home microbiome study in southern New England revealed that microbial richness was associated with the presence of pets, water leaks and suburban locations, and that microbial community variation was associated with air conditioner usage and occupancy. Additionally, the quantity of specific bacteria, such as *Lactobacillus johnsonii*, was reported to be associated with occupant density [18]. The results showed that microbial α- and β-diversity were associated with different factors in homes, consistent with our results. A study of a local university housing facilities showed that season was an important factor shaping microbial α- and β-diversity but was not associated with indoor fungal quantity. Even a short geographic distance was important in shaping fungal community variation [47]. There have also been several continental-scale microbiome studies of home environments, revealing the importance of geographic distance and outdoor factors in structuring microbial communities, especially fungal communities, in the indoor environment [17, 48]. However, these studies did not conduct qPCR quantification; thus, the associations between environmental factors and microbial quantity were not clear. The major finding of our study is the concordant/discordant pattern for factors associated with α-diversity, β-diversity and quantity, which was not discussed in previous studies. It is possible that environmental factors, such as low latitude or high relative humidity, serve as suitable ecological niches for the proliferation of certain microbial species, leading to changes in total microbial quantity and the microbial community. This hypothesis is supported by a laboratory chamber study on cut carpets from homes. The study showed that elevated humidity led to the growth of a limited number of fungal taxa, including *Aspergillus* and *Penicillium* [49]. Thus, the variation in the hotel microbiome may be similar to that in the chamber model in the laboratory, but with more impacting factors, such as geographic location, further influencing the source of fungi. Overall, more continental-scale indoor microbiome studies are needed to test if the model reported in our study is applicable in other environmental settings.

### High abundance of bacterial and fungal taxa in hotel rooms

Most of the high-abundance bacteria in hotel rooms are ubiquitous taxa in outdoor environments and are widespread in a wide range of habitats, such as *Ralstonia*, which is commonly found in soils, rivers and lakes, and *Pelomonas*, which is found in water and soil [50]. Thus, it is common to find these taxa in hotel rooms, but due to their ubiquitous nature, these bacteria are frequenctly reported as sources of contamination for reagents and laboratories that impact the accuracy of sequence-based microbiome analyses [51]. For the hotel samples, the negative control in the amplification process indicated no DNA or microbial contamination. We also checked the sequencing results from other samples in the same MiSeq run as the hotel samples and found that most samples did not contain *Ralstonia* or *Pelomonas* species (data not shown), confirming that they are not general contaminants in the sequencing laboratory. A previous study reported that human-associated microbes, especially those on skin, were an important source of the indoor microbiome, accounting for 4% to 40% of the total bacterial load [28]. In the hotel dust samples, the abundance of human-associated microbes, such as *Acinetobacter* (4.5%), *Propionibacterium* (3.2%), *Corynebacterium* (1.4%), *Streptococcus* (0.25%), *Staphylococcus* (0.33%), *Bifidobacteria* (0.21) and *Kocuria* (0.41%), was lower than that of the environmental taxa. The total contribution of human-associated taxa in hotels was similar to some previous studies in home residences [28].

In hotel rooms, *Aspergillus* was the dominant fungus, accounting for on average one fourth of the total fungal load. The negative control and other samples in the same run also confirmed that the taxon was not from laboratory contamination or technical bias. The abundance of *Aspergillus* was significantly higher in low-latitude hotels (low-latitude mean: 46.3%, middle-latitude mean: 18.8%, high-latitude mean: 9.3%), and in seven low-latitude hotels, *Aspergillus* even accounted for 80% of the total fungal load. This result was consistent with qPCR quantification results for primers targeting *Aspergillus* and *Penicillium*, which accounted for 53.2%, 30.1% and 10.7% of total fungi in low-, middle- and high-latitude hotels, respectively (Table S7). In addition, a previous study of 1,200 homes across the US showed that the contribution of *Aspergillus* can vary dramatically from 0% to >95%, even with a mean proportion of only ∼4% [17], indicating that it is possible to detect variable amounts of *Aspergillus* in indoor environments. In addition to latitude, proximity to the sea and visible mold were also associated with a high abundance of *Aspergillus*.

These environmental characteristics are all correlated with indoor air moisture and relative humidity (RH), which are key factors regulating fungal growth the in laboratory. Incubation of room dust at an 84-86% RH resulted in a 45-fold increase in *Aspergillus* and *Penicillium* [52]. A chamber study demonstrated that *Aspergillus* and *Wallemia* growth occurred at >80% RH from carpet after one week of incubation [49]. It is worth noting that some commonly reported airborne fungal genera [10, 48] are not at a high abundance in hotels, such as *Penicillium* (2.2%), *Cladosporium* (0.26%), *Acremonium* (0.44%), *Alternaria* (0.06%), *Fusarium* (0.08%), *Mucor* (<0.001%), *Stachybotrys* (0.12%), *Trichoderma* (0.32%) and *Trichophyton* (<0.001%). In contrast to *Aspergillus, Penicillium* was more abundant at the middle and high latitudes (3.6% and 2.5% vs 0.7%, p < 0.001). Indeed, previous studies identified *Penicillium* as one of the major airborne fungi in buildings that were mainly sampled in middle- or high-latitude countries, such as Norway, France and Poland [53-55]. Global-scale sampling of settled dust revealed that *Penicillium* was present at a low abundance in low-latitude indoor environments [48], consistent with our observations in hotels.

From a health perspective, *Aspergillus* is abundant in hotel rooms, and the species can produce fungal fragments, such as pieces of spores or hypha, microbial volatile organic compounds or mycotoxins, and lead to various allergic or inflammatory symptoms in occupants, such as cough, wheezing and headaches [10, 56, 57]. Thus, it may be necessary to routinely monitor the mold growth and *Aspergillus* quantity in hotel rooms, especially at low latitudes.

### Factors associated with microbial richness in hotel rooms

The latitudinal diversity gradient (LDG) theory states that biodiversity declines with latitude, and the theory is supported by the majority of ecological studies [58, 59]. However, deviation from this pattern has also been reported [60, 61]. An indoor study also reported higher fungal diversity in temperate zones than in the tropics [48]. Hillebrand conducted a meta-analysis of 600 studies and found that the strength of the diversity gradient increased greatly with organismal body mass, possibly due to energy use and dispersal limitation [58]. Thus, microbial richness is less affected by latitude than the richness of larger vertebrates. In this study, we found that latitude did not affect the microbial richness in hotel rooms, which is consistent with Hillebrand’s prediction. A relationship between mechanical ventilation and fungal quantity indoors has been reported in many studies. Mechanical ventilation equipped with good filters can remove coarse airborne particles from outdoors, which was reported to reduce the indoor fungal concentration [10, 62]. However, improperly maintained ventilation systems could also act as sources of contamination and increased the fungal concentration [63]. Mechanical ventilation was also reported to change the bacterial community composition [64], but in this study, we found that it reduced bacterial richness but did not alter community composition. Thus, the role of mechanical ventilation is complex and may vary for different ventilation systems and building designs. We also found that wall-to-wall carpet reduced the fungal richness in hotels. Most previous studies reported that carpet was a sink for fungi that increased microbial diversity [65], although contradictory evidence is also available and suggest that the effect of carpet may depend on the actual carpet type [10]. Another possible explanation is that the daily routine cleaning of carpets reduces the fungal richness in hotels.

### Factors structuring microbial composition variation

In addition to latitude, seaside location and visible mold were associated with fungal β-diversity and quantity, and geographic distance was also an important factor determining microbial composition. A previous study showed that community similarity was negatively correlated with geographic distance for both bacteria and fungi, but this relationship was stronger for fungi [20]. We confirmed the pattern by showing that most indoor fungal OTUs were locally distributed, whereas bacterial OTUs were more widely distributed. This is because the source of indoor fungi is mainly from outdoor fungi, which is more geographically patterned than the outdoor bacteria [20]. In addition, most fungal species in indoor environments are not from human sources [66], thus, they cannot be distributed through human travel and movement, which further limits fungal distribution ranges.

It has been suggested that the indoor fungal composition is primarily shaped by global rather than indoor factors. A survey of indoor environments (n=72) revealed that global factors, rather than building design and materials, determine indoor fungal composition, and the indoor fungal assemblage represents a subsample of the outdoor fungal community [48]. Most fungi may not grow or proliferate in the indoor environment; thus, the indoor environment mainly serves as a passive collector for the outdoor fungal biome [47, 48]. However, a recent survey of university residences in California found that fungal composition clustered by indoor surface type, suggesting that some fungal species do grow or adhere to certain surface types, which leads to composition variation [66]. In this study, we found that global factors, such as latitude, were the most important factors shaping the microbial community, and indoor factors, such as the quality of the interior and floor surface type, were less important. The indoor factors were significantly associated with bacterial composition variation but only marginally associated with fungal composition, suggesting that they were more involved in structuring the bacterial community than in structuring the fungal community.

### Characteristics not associated with microbial diversity and variation

Urban/rural location was reported to be associated with microbial diversity or quantity [10]. Farming environments have more diverse fungal resources than urban areas, which leads to a reduction in early childhood asthma in rural areas [13]. In this study, we did not detect an association between urban/rural locations and microbiomes. This could be due to the homogenization of the microbiome in hotel rooms caused by daily routine cleaning or because urban/rural location was correlated with other strongly influential factors, such as seaside location (ρ=-0.54, p < 0.001, Pearson’s correlation) and latitude (ρ=0.35, p = 0.003). Other hotel characteristics, such as the number of stars, building age and size of the hotel, were not associated with microbial diversity and composition. Thus, factors associated with high-ranking hotels do not change the microbiome community in rooms.

### Study strength and limitations

The strength of this study is that it is the first hotel microbiome study spanning large continental regions in Asia and Europe, providing useful resources for future indoor or hotel microbiome studies. The environmental metadata of the study were collected by a professional hygienist; thus, characteristics such as visible mold or dampness were assessed by professional standards, making them more consistent than the self-reported observations of residents in some home studies.

One limitation of this study is that sample from only one site were collected in each hotel room. Previous studies showed that sampling surface type and indoor location affected microbial community composition [16, 67]; thus, sampling dust at multiple sites can lead to a more comprehensive assessment of indoor microbiome composition. As the floor surfaces in hotels are frequently vacuum cleaned, other sites such as the top side of the mirror cabinet in the bathroom or curtain surfaces may be appropriate sampling locations for future studies. In this study, dust was collected by cotton swabs with a swabbing area of 1 x 60 cm for each sample; thus, the quantitative estimates were presented as the number of fungi per square meter. Since the hand pressure for swabbing may vary across the sampling sites, biases can be introduced in the sampling process, and the quantity results should be interpreted with caution. However, as the dust swabs were all performed by a single hygienist, the variation in the sampling process should be minimal. In addition, although we screened negative controls and other sequencing results on the same MiSeq run to show that technical biases and laboratory contamination were not major factors shaping the microbiome variation, these factors still may have affected our results to a certain extent.

## Conclusions

We presented here the first continental-scale hotel microbiome study spanning 19 countries and revealed the detailed microbial composition and features in this environment. This is the first study to show that the environmental factors associated with microbial β-diversity are also associated with absolute quantity but not associated with α-diversity. *Aspergillus* was the most abundant fungus in hotel rooms and was negatively associated with latitude, whereas *Penicillium* was much less abundant especially in low-latitude hotels. Most bacteria in hotel rooms were ubiquitous species sourced from outdoor environments instead of from human sources (10-15%). We uploaded all the data to QIITA platform to facilitate research progresses on the built environment. In the long term, these data can be integrated into a larger meta-analysis study to study human microbial exposure or identify a “healthy building microbiome” to promote human well-being in a general indoor environment.

## Materials and Methods

### Sample and data collection

Dust swab samples were collected by a professional hygienist from 68 hotels in 19 European and Asian countries from October 2007 to May 2009. We collected dust samples in one room in each hotel by using cotton swabs to swab the upper half of the doorframe. Two samples were collected in each room with a swabbing area of 30 cm^2^ (1 x 30 cm) on the left- and right-hand sides of the doorframe. Each swab was rotated slowly and moved back and forth 3 times over the surface. The swabs were stored in a −80°C freezer after sampling.

More than 30 outdoor and indoor hotel characteristics were collected for each sampling site. We conducted Pearson’s correlation analysis of all the characteristics to reduce correlated redundant characteristics (ρ > 0.7). For example, annual precipitation was highly correlated with latitude (ρ = −0.82); thus, only latitude was kept for further analysis. In total, 16 environmental factors were included in further analyses, namely, latitude, surrounding traffic (heavy or light traffic), distance to an airport, proximity to the sea, location of the sampling site (rural, suburban, urban, or megacity), number of stars of the hotel, building age of the hotel, redecoration age of the hotel, size of the hotel, quality of the interior, floor level, visible mold, mechanical ventilation, floor surface type in the hotel room, air conditioner in the wall and odor in the hotel room. The detailed grouping of variables is presented in Table 1. For details of the sampling process, please refer to a previous publication [34].

### Microbial DNA extraction and amplicon sequencing

Total genomic DNA was extracted by an E.Z.N.A. Soil DNA Kit D5625-01 (Omega Bio-Tek, Inc., Norcross, GA, USA), which uses bead beating and spin filter technology to extract DNA. Total fungal DNA was extracted by Fast DNA SPIN extraction kits (MP Biomedicals, Santa Ana, CA, USA). The quality and quatity of extracted DNA were evaluated by agarose gel electrophoresis, a NanoDrop ND-1000 spectrophotometer (Thermo Fisher Scientific, Waltham, MA, USA) and a microplate reader (BioTek, FLx800), and all 68 samples passed the quality-control step and thus qualified for amplicon sequencing. The library was prepared by a TruSeq Nano DNA LT Library Prep Kit from Illumina. The forward primer 338F (ACTCCTACGGGAGGCAGCA) and reverse primer 806R (GGACTACHVGGGTWTCTAAT) were used for bacterial 16S rRNA gene V3-V4 region amplification, and the amplification region was 480 bp in length. The forward primer ITS5 (GGAAGTAAAAGTCGTAACAAGG) and reverse primer ITS2 (GCTGCGTTCTTCATCGATGC) were used for fungal ITS1 region amplification, and the amplification region length was 250 bp. Sample-specific 7-bp barcode sequences were incorporated into primers for multiplex sequencing. Before sequencing, the library was evaluated by an Agilent Bioanalyzer and a Promega QuantiFluor with a Quant-iT dsDNA Assay Kit. Paired-end sequencing was performed according to the manufacturer’s instructions with the Illumina MiSeq platform and a MiSeq Reagent Kit v3 (600 cycles) at Shanghai Personal Biotechnology Co., Ltd (Shanghai, China).

### Bioinformatics and sequence analysis

The raw sequences with a short length (< 150 bp), low Phred score (< 20) and ambiguous bases and mononucleotide repeats longer than 8 bp were removed [68]. Raw sequences were extracted according to the barcode sequence and assigned to the respective samples. Flash (v1.2.7) was used to assemble the paired-end reads with a minimum overlap between forward and reverse reads > 10 bp and no mismatches [69]. Many of the following analyses were conducted with the Quantitative Insights Into Microbial Ecology (QIIME, v1.8.0) platform [70] and R packages. Chimeric sequences were removed by USEARCH (v5.2.236) [71]. It has been shown that the erroneous reads from PCR in the amplicon preparation step and sequencing error lead to the overestimation of microbial diversity [72]. Thus, we conducted a stringent quality-filtering step to extract high-quality data and set the operational taxonomic unit threshold (c value) to 0.01% in QIIME and other parameters following a previous suggestion [72]. The remaining high quality sequences were clustered into operational taxonomic units (OTUs) with 97% sequence identity. A representative sequence was picked for each OTU and blasted against the Silva database [73] for bacteria and UNITE database [74] for fungi to obtain taxonomic classification information. Rounded rarefied analysis was conducted to standardize the sequencing depth among all samples. The α-diversity index of the observed species was calculated based on the OTU table. The β-diversity was represented by principal coordinate analysis (PCoA) [75] based on UniFrac distance metrics [41]. Linear regression and a Kruskal-Wallis test were performed by IBM SPSS Statistics (v21.0). Permutational bivariate and multivariate analysis of variance (Adonis) [76] was conducted by the vegan package in R, with a Bray-Curtis dissimilarity distance matrix and 10,000 permutations. The multivariate analysis was calculated with a forward stepwise approach. The factor with the lowest p value in the bivariate analysis was input first, and the factor with the second lowest p value was input second, and so on (inclusion level p = 0.2).

## Declarations

### Ethics approval and consent to participate

Not applicable.

### Consent for publication

Not applicable.

### Availability of data and material

Sequencing data was deposited in Qiita with study ID 12274 (https://qiita.ucsd.edu/study/description/12274).

### Authors’ contributions

DN and GC collected dust samples in hotels. DN and YS designed the project. XF, YL, QY and YS carried out the analysis. XF, YD, XZ, DN and YS drafted the manuscript. All authors read and approved the final manuscript.

## Acknowledgments

We thank Personalbio (www.personalbio.cn) for help in sequencing analysis.

